# Cell Type Specific Responses of the Endoplasmic Reticulum Dynamics to Environmental Stress

**DOI:** 10.1101/2024.01.23.576814

**Authors:** Yiheng Zhang, Jiazheng Liu, Zhengzhe Sun, Jinyue Guo, Jingbin Yuan, Yajun Xue, Tianbao Qiu, Pei Wang, Benfeng Yin, Honglan Yang, Liting Zhai, Hua Han, Xiaojuan Li

## Abstract

To overcome the challenge of balancing imaging speecd and resolution, which currently limits the accurate identification of structural and dynamic changes in the study of endoplasmic reticulum (ER) in plant cells. This research employs structured illumination microscopy techniques to achieve super-resolution real-time imaging of the ER in live Arabidopsis materials. Additionally, a self-supervised denoising framework (Blind2Unblind) was optimized to further enhance the signal-to-noise ratio of rapid microscopic imaging. Based on the images with high quality, a method for quantitative analysis of ER structures using time-lapse images is developed. Moreover, detections of changes in ER structures under environmental stress are conducted to verify the effectiveness of the method. Moreover, correlation analyses of various parameters indicate a significant positive correlation between the area and length of tubular ER with the number of growth tips and tricellular junctions, while the area of ER cisternae and bulk flow exhibits a significant negative correlation with the area and length of tubules. The super-resolution imaging and dynamic analysis method developed in this study will provide new technical approaches for further elucidating the function and regulatory mechanisms of the plant ER.

## Introduction

The endoplasmic reticulum (ER) is a continuous membrane system forming flattened sacs within eukaryotic cell cytoplasm, involved in numerous cellular physiological activities. Being highly dynamic, the movement and remodeling of the ER are intricate, occurring across a broad spectrum of scales (Griffing et al., 2017; Kriechbaumer et al., 2018). This high mobility is associated with its structural components, such as individual proteins, protein complexes, and unique fusiform bodies found in Arabidopsis, within the dynamics of ER particles. ER particle dynamics involve the movement of these components through the ER’s lumen or membrane, contributing significantly to its high dynamism (Hawes et al., 2001; Pain and Kriechbaumer, 2020). ER remodeling, involving changes in the tubular network, cisternae, and stable points, is propelled by the actomyosin cytoskeleton, displaying an approximate streaming speed of ∼3.5 μm/sec within the inner plane of the ER (Ueda et al., 2010; Pain and Kriechbaumer, 2020). An interactive relationship exists between ER particle dynamics and ER remodeling. For instance, the topological arrangement of the ER tubular network corresponds to the movement of luminal particles that diffusive component prefer to domain in the tubular junctions and the fast flow component distribute in tubules (Holcman et al., 2018) and when the ER accumulates new palmitate metabolites, it exhibits a lamellar structure and undergoes rapid growth (Shen et al., 2017).

Previous studies have focused significantly on the intricate relationship between the structure and function of ER. ER-Golgi intermediate compartment (ERGIC), is rich in tubules and vesicles which involve in lipid convey through organelle contacts or secretory vesicles when lipid get into ERGIC (Appenzeller-Herzog and Hauri, 2006; Fagone and Jackowski, 2009). In addition, depolarization of t-tubule membranes can lead to conformational changes in voltagedependent Ca2+ channels which related to calcium signaling (Fill and Copello, 2002). Ribosomes have high density on the cytosolic surface of cisternae, which makes cisternae be the main site of synthesis, folding and post-translational modifications for secreted or membrane-bound proteins (Schwarz and Blower, 2016). Due to its role in synthesizing proteins and lipids, the structure and dynamics of the ER exhibit variations across different cell types. ER structure within specialized cells, like pancreatic secretory cells and B cells, primarily comprises sheet-like formations. Cells engaged in lipid synthesis, calcium signaling, and acting as contact points for other organelles possess an ER primarily made up of tubules. Those tight relationship between structure and function display in different type of cells. For instance, adrenal, liver, and muscle cells exemplify specialized cells primarily characterized by a tubular network which reflecting those specific functions (Baumann and Walz, 2001; Schwarz and Blower, 2016). As for plants, they notably harbor a unique architectural feature termed the “desmotubule,” originating from the ER, aiding in the creation of channels that stretch from the ER to the cell wall. These channels seamlessly interconnect the cytoplasm of neighboring cells (Wright et al., 2007; Knox et al., 2015; Nicolas et al., 2017). In moss, during bud formation, the ER initially takes on a condensed network of membranes that later transforms into a complex and intricately reticulated configuration as the cells grow. Cisternae, accompanied by their attached ribosomes, primarily serve as sites for protein synthesis, while tubules are commonly associated with lipid synthesis (Voeltz et al., 2002; Shibata et al., 2006). Peripheral ER, referring to ER outside the nuclear envelope, exhibits distinct characteristics between yeast and higher eukaryotes. In yeast, it exhibits a closer distribution to the cell cortex, with only a few tubules extending to the nuclear envelope. Conversely, in higher eukaryotes, peripheral ER extends across the entire cytoplasmic volume (Shibata et al., 2006). Cortical ER, localized near the plasma membrane, presents as a consistent feature in yeast cells, linked to the coat protein complex I (Lavieu et al., 2010), but the cortical ER is usually found in specific cell types, like muscle cells, neurons and other specific cells. Throughout various stages of the cell cycle, significant diversity exists in the extent of spatial reorganization. Additionally, the transition from sheet-like structures to tubules varies among different mammalian cell lines especially in mototic cells (Puhka et al., 2012).

Instead of assigning individual sensing or response mechanisms to each organelle, cells prefer to achieve cellular efficiency by utilizing the ER as the primary sensor and responder during periods of stress. This function is demonstrated during periods of nutrient scarcity when cells elevate the levels of CLIMP63 and p180, proteins associated with microtubules, to regulate the movement of both the ER and lysosomes, facilitating effective autophagic degradation and appropriate reformation (Zheng et al., 2022). Regarding proteins situated in the ER lumen, SARS-CoV-2 ORF8 evades degradation within the host cell, inducing ER stress and leading to the abnormal morphogenesis of the ER characterized by tangled bends, potentially resulting in heightened protein synthesis (Liu et al., 2022b). Membrane contact sites are a direct link between organelles, providing a non-vesicular, direct and rapid means of exchanging material information for communication between organelles. Thus, during cellular stress, the ER can also interact with other organelles to respond. ER-mitochondrial contact sites (EMCS) play a significant role in physiological processes and mitochondrial functions such as Ca2+ transfer, lipid metabolism, autophagy, mitochondrial dynamics, and morphogenesis (Kornmann et al., 2009; Friedman et al., 2011; Hamasaki et al., 2013). During stress, the proteins within EMCS can regulate interactions between the ER and mitochondria, influencing mitochondrial autophagy (Li et al., 2022). ER–autophagosomal membrane contact site (EACS) can regulate plant autophagy with the lipid-binding protein ORP2A locates in EACS (Ye et al., 2022).

The accuracy and resolution of electron microscopy have rapidly improved, allowing for more nuanced combinations and variations (Davidowitz et al., 1975; Puhka et al., 2012). Prior to electron microscopy, research on the neuronal endoplasmic reticulum relied heavily on the enduringly popular Nissl stain, which has limitations and drawbacks. It fails to reveal details about specific cellular structures or functions beyond the rough endoplasmic reticulum, has limited utility for living tissue, and cannot visualize fine structures (Goetz, 2013). Because the time scale of ER changing is in sec, neither of these methods could visualize the dynamic changes of the endoplasmic reticulum in living cells. Since the initial visualization of the ER network using fluorescent dyes (Quader and Schnepf, 1986), the use of fluorescent dyes and techniques for living cell imaging has rapidly progressed. To evaluate ER dynamics, most methods relied on standard fluorophore bleaching/photoactivation experiments to track the recovery of bleached or photoactivated fluorophores in the ER network (Sparkes et al., 2009; Griffing et al., 2017; Holcman et al., 2018). Confocal microscopy, or combined with photoconversion pulse-chase, has been widely used for high-resolution imaging in living cells to visualize the ER structure (Holcman et al., 2018; Pain and Kriechbaumer, 2020). Additionally, Lu et al., (2023) employed Structured Illumination Microscopy SIM to image ER structures in mammalian cells, enhancing the resolution from high to super-resolution. Over time, live-cell microscopy techniques, such as live-cell STED combined with in situ cryo–electron tomography, have observed ribosome-associated vesicles primarily located in the cell periphery, a common feature across various cells and species (Carter et al., 2020). The majority of present studies focus on qualitatively describing the structure of plant ER, while quantitative analysis of its structure and dynamic changes remains inadequate. Obtaining high-resolution cellular structures relies on imaging tools with high spatial resolution. However, these tools often compromise imaging speed, which is crucial for fast-changing biostructures. Previous studies have failed to strike a balance between these aspects, resulting in deficiencies in quantitative analysis based on high/super-resolution imaging. In summary, the fast and stable acquisition of high-resolution image data with a high signal-to-noise ratio poses a significant challenge, limiting the accurate identification of large-scale cellular structures. Additionally, a notable limitation in previous studies is their focus on quantifying ER structural information in a single cell type (Pain et al., 2019; Lu et al., 2023). Neglecting variations in ER structure across different cell types within the same organism.

RALFs (Rapid alkalinization factors) is a class of multifunctional plant cytokines which is a polypeptide with 5 kDa and induce a rapid increase in extracellular pH in plants (Pearce et al., 2001). Recent studies have revealed that RALF plays an improtant role in biological process. For instance, RALF22, RALF17 and RALF23 can be involoed in plant immune responses, with the regulation involved with RALF23, FERONIA (FER) can acts as a RALF-regulated scaffold in receptor kinase complex assembly (Stegmann et al., 2017; Franck et al., 2018; He et al., 2023). Meanwhile, RALF1 and RALF23 bind to the CrRLK1L receptor FER promote root, leave and seedling growth (Franck et al., 2018; Pacheco and Estevez, 2022; Song et al., 2022). RALF34 interacts with the THESEUS 1 (THE1) receptor which regulated lateral root initiation and defense responses(Gonneau et al., 2018). The pollen tube growth, polytubey block, and cell-wall integrity also contain the members of RALF family with other receptor like FERONIA (FER), ANJEA (ANJ), HERCULES RECEPTOR KINASE 1 (HERK1) receptor-like kinases ANX1/2 (ANXUR 1/2) and BUPS1/2 (Buddha’s Paper Seal 1/2) (Mecchia et al., 2017; Feng et al., 2019; Ge et al., 2019; Zhong et al., 2022). Some nematode-encoded RALF-like peptide also play the typical activities in parasitism of plants (Zhang et al., 2020). However, almost studies are focus on the physiological function instead of its physical feature. Plant extracellular pH is acidic (pH 5.7), and many physiological and external environmental factors cause changes in extracellular pH (Liu et al., 2022a). The basic alkali feature of RALF has been ignored for a long time.

We utilized super-resolution imaging using SIM, known for its low phototoxicity, non-damaging nature, and low background noise detection, to visualize the ER structure. After acquiring images with precise ground truth values, we utilized the improved denoising framework (BlindU2blind) specifically designed for SIM data to enhance image quality. Introducing a new architecture, Swint-ResU-Net, a U-shaped network that integrates Swin Transformer modules and residual modules, aimed at handling intricate biological tissue data for the automatic segmentation of endoplasmic reticulum structures. Utilizing the segmentation results from Swint ResU-Net, we constructed a tubule network, and identified stable points within the network, including growth tip, three-way junction and multi-way junction. We quantitatively analyzed the variations in ER structure across three distinct parts of Arabidopsis thaliana subjected to abiotic stress at different growth stages. Our findings revealed that-----. Leveraging the exceptional performance of our segmentation model and ER structure quantification, we introduced innovative approaches for quantifying ER structures in plants. Furthermore, we unveiled distinctions between plant segments under stress, offering valuable insights into endoplasmic reticulum structural changes.

## RESULT

### Segmentation of ER structure

The cotyledon, hypocoty, and elongation zone of Arabidopsis thaliana were the three growth sections whose ER we examined (Figure1.A). We obtained the super-resolution structure of those growth parts in growth days 3-5 using the SIM pictures (Figure 1B-D). Coupled with our earlier research (Wang et al., 2023) preliminarily established an implicit lossless self-supervised denoising framework-Blind2Unblind. Based on SIM data, we refined this framework, which addressed pixel-independent noise, resolved input loss, and went on to handle structured noise reduction (Figure 1E–G; Supplemental Figure 1).

**Figure 1.**
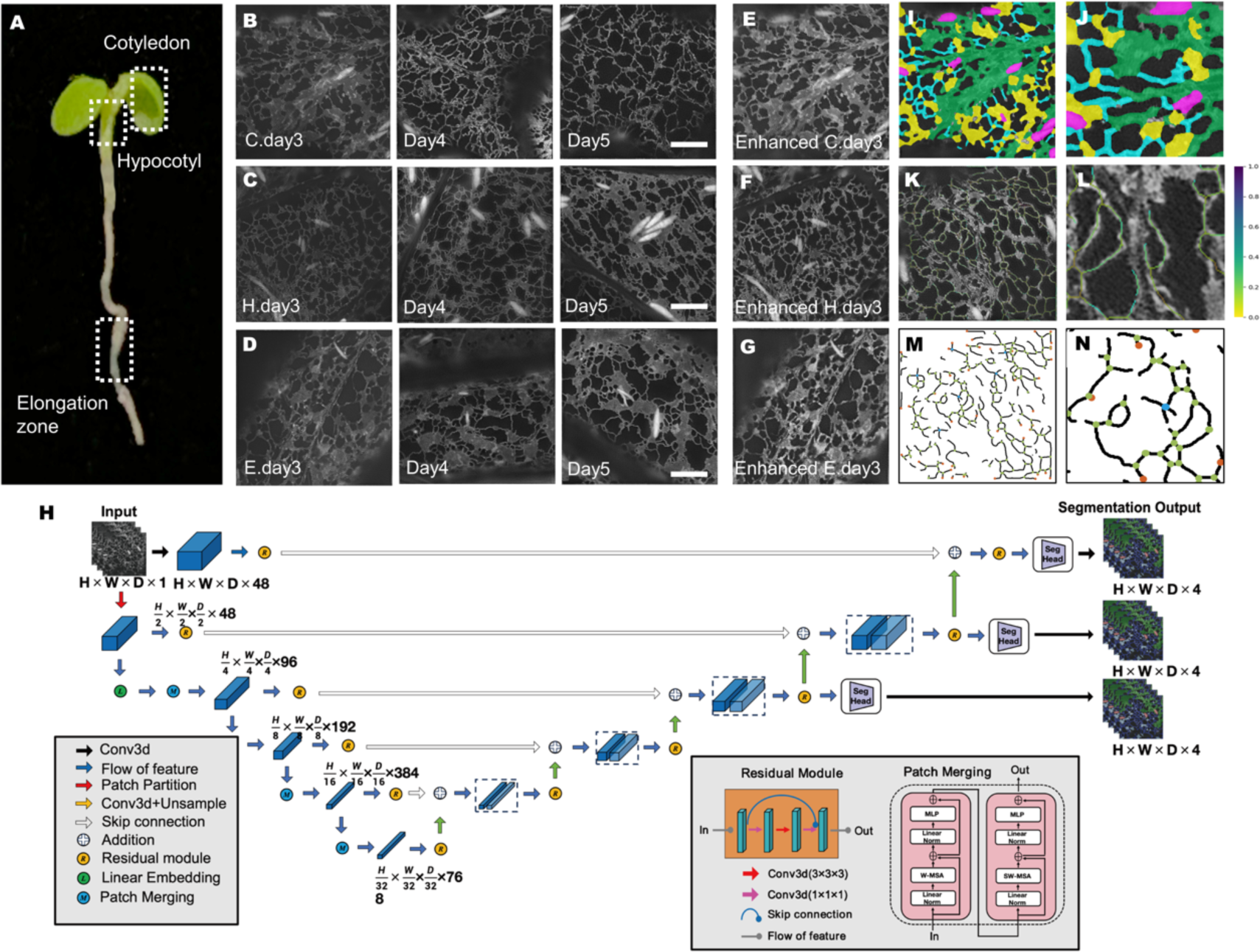
Segmentation and identification of ER structure. **(A)** Cells from three growth regions of Arabidopsis were imaged: cotyledon, hypocotyl, and elongation zone. **(B)-(D)** ER structures from cotyledon, hypocotyl, and elongation zone at growth days 3, 4, and 5, imaged using SIM and enhanced by Blind2Unblind. Scale bar = 2μm. **(E)-(G)** Enhanced images of ER structure of cotyledon, hypocotyl, and elongation zone in day3 by BlindUn2blind. **(H)** Overview of the Swint-ResU-Net architecture. The input to our model is Time Series SIM images with single channels. The Swint-ResU-Net creates non-overlapping patches of the input data and uses a patch partition layer to create windows with a desired size for computing the self-attention. The encoded feature representations in the Swin transformer are fed to a Res-decoder via skip connection at multiple resolutions. Seg Head denotes the final segmentation layer. Final segmentation output consists of 4 output channels corresponding to cisternae, tubules, flow and fusiform bodies. **(I)-(J)** Comprehensive segmentation results highlighting four distinct parts of the ER structure marked by different colors: Tubule (cyan), cisternae (bright yellow), bulk flow (green), and fusiform body (magenta); **(J)** Detailed enlarged display of **(I)** **(K)-(L)** Extraction of tubule information of ER. The colors on the tubule present the width of the tubule; **(L)** Detailed enlarged display of **(K)** **(M)-(N)** Identification of nodes in ER structur, including growth tips (orange), three-way junctions (light green), and multi-way junctions (light blue); **(N)** Detailed enlarged display of **(M)**.

Manual annotation methods cannot meet the efficiency needs of large-scale reconstruction, while neural networks have shown impressive performance in image segmentation. Currently, the base model of U-Net and its variants are still widely used in medical and biological image segmentation, achieving satisfactory segmentation results. Recently, transformer-based models have emerged as a superior choice across various domains such as natural language processing and computer vision (Dufter et al., 2021; Liu et al., 2021; Hatamizadeh et al., 2022a; Hatamizadeh et al., 2022b).

We introduce a pioneering network architecture, named Swint-ResU-Net, amalgamating Swin Transformer modules and residual modules to tackle the complexities within biological tissue data. This fusion aims to enable the automatic segmentation of endoplasmic reticulum structures. The architecture employs 3D Swin Transformer modules in its encoding phase for downsampling. This process progressively reveals environmental information while expanding the network’s receptive field. In contrast, the decoding structure relies on convolutional neural networks (CNN), specifically utilizing ResNet. This choice facilitates gradual upsampling, ensuring the restoration of image accuracy. Moreover, it establishes communication of contextual information via skip connections operating at various resolutions. The empirical results underscore the criticality of a substantial receptive field, effective contextual information exchange, and a deep network structure in accomplishing ER segmentation tasks (see Figure 1E and Supplemental Figure).

Utilizing Swint-ResU-Net segmentation, we discerned distinct regions within the ER based on their morphological traits. Tubules and cisternae, the primary ER components, demonstrate ER remodeling. They are propelled by the actin-myosin cytoskeleton, leading to a bulk translocation known as bulk flow, inducing ER streaming. Additionally, we identified a fusiform body within the image (Figure 1I,1J). Tubules assume a crucial role within the ER network. Consequently, we conducted an extraction of tubule-related information and proceeded with an analysis focusing on the width of the tubule network (Figure 1K,1L). Regarding stable points, specifically stable regions surrounding ER remodeling sites, their association extends to both cisternae and tubules. Our identification process involved nodes within the tubule network, encompassing growth tips, three-way junctions, and multi-way junctions (Figure 1M,1N). To validate the accuracy of our segmentation, we accessed ground truth data for comparison, conducting a pixel-by-pixel evaluation. The resulting ground truth test underscored the pixel accuracy of Swint-ResU-Net segmentation, meeting the anticipated standards. The accuracy of ER structure ranged between 94% and 99% in comparison to the ground data, exhibiting remarkable precision across different ER components (tubule: 94.7%, cisternae: 95.62%, bulk flow: 94.56%, fusiform body: 99.49%) (Supplemental Figure). The identification of three-way junctions relied on the tubule network configuration, specifically at points where three tubules converge into a single node. Subsequently, leveraging the segmentation results pertaining to structural parameters, we performed a comparative analysis between the identified three-way junctions (tubule identification results) and their respective ground truth data (Supplemental Figure). The accuracy of three-way junction was 95.12% and it was 92.13% in the identification of growth tip.

### Dynamic changing of ER structure

As a highly dynamic organelle, ER undergoes continuous changes in its network connections, involving the movement of thousands of tubules and cisternae, as well as the rapid flow of bulk material. To scrutinize these dynamic alterations in the ER’s structure, the initial frame (1st frame) of the ER time series imaging was superimposed onto the 20th frame image (Figure 2A). The analysis revealed minimal structural changes in the cisternae and bulk flow. However, the alignment of the tubular ER’s red and green components displayed notable disparity, indicating significant alterations (Figure 2B). Simultaneously, the time-series dynamics maps exhibited discernible morphological changes in various ER structures, including the growth tip and multi-way junction (Figure 2C), three-way junction (Figure 2D), cisternae (Figure 2E), and tubule (Figure 2F).

**Figure 2.**
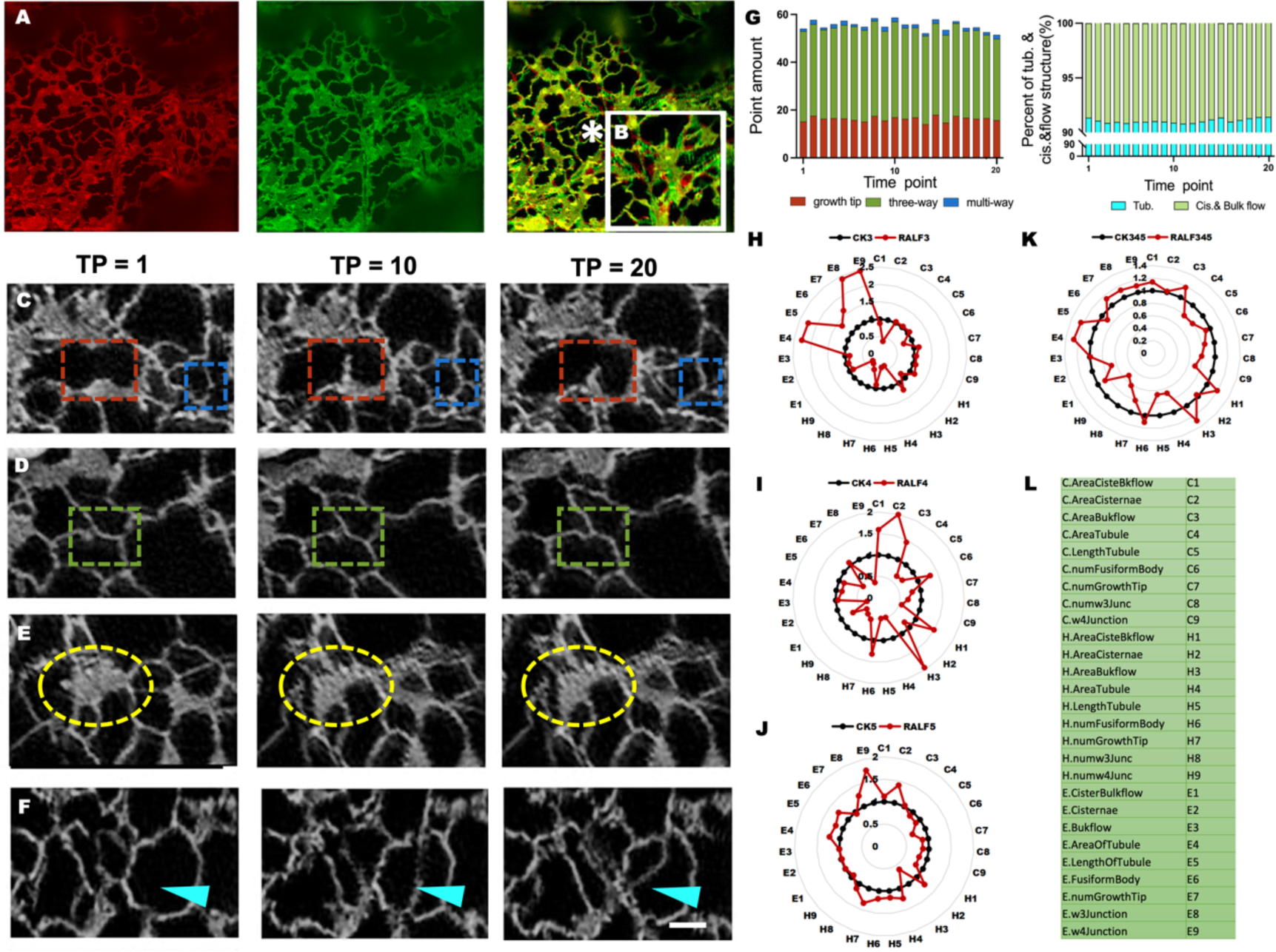
Dynamic changing of ER structure. **(A)-(B)** ER structure images of cotyledon in day 3 at the time point was1 (red), 20 (green) and merged in yellow. **(B)** Enlarged display of details in the asterisk area. **(C)-(F)** Structure changing in continuous time-point (1-20). **(C)** Changing of growth tip (red box) and multi-way junction (bule box); **(D)** Changing of three-way junction (green box); **(E)** Changing of cisternae (bright yellow ellipses); **(F)** Changing of tubule (cyan arrows indicate areas). **(G)** Quantitative analysis of the ER shown in a. Left, quantification of point amount changing in time point, including growth tip, three-way junction, and multi-way junction. Right, percentage of areal tubule(cyan) and areal cisternae and bulk flow (mint green) in time point (30ms per time point). **(H)-(J)** Dynamics of nine ER structural parameters under CK and RALF treatments at day3,4 and 5. **(K)** Comprehensive analysis of the dynamics of nine structural parameters of the ER on days 3, 4 and 5. **(L)** Table of abbreviated form of nine ER structural parameters used in **(H)-(K)**.

SIM offers high spatial-temporal resolution of ER structures while minimizing phototoxicity, preventing damage, and reducing background noise, making it ideal for live cell imaging. Employing a clear segmentation of the ER structure within our network, we conducted quantitative analysis of nodes—encompassing growth tips, three-way junctions, and multiway junctions (Figure 2G, left)—as well as the proportional area occupied by tubules, cisternae, and bulk flow within the ER (Figure 2G, right) across the entire ER structure’s field of view. The quantification of morphological features at various time points within the branched network (nodes and tubules) and the flat membrane network (cisternae and bulk flow) serves as additional evidence corroborating the highly dynamic nature of the ER.

ER stress induces morphological changes in the ER structure. The protein GFPase Rab7, known for modulating lysosome-ER contact sites to regulate ER shape, exhibits an inhibitory effect when its expression is suppressed, resulting in the enlargement of sheet-like ER structures that extend towards the cell periphery (Mateus et al., 2018). Moreover, environmental stressors, such as heat shock, can diminish the tubular ER network and prompt an ER stress response characterized by an increase in cisternae size (Pain et al., 2019). Our quantitative data analysis revealed that rapid pH changes in the environment, triggered by RALF, caused distinct morphological alterations in the ER structures across various growth stages. Following RALF treatment on the 3rd day seedling, notable changes were observed in the elongation zone, where the number of multi-way junctions increased significantly by 145.05% compared to the control (CK). Simultaneously, the region exhibiting the most substantial reduction (72.16%) in trigeminal points was also situated within the elongation zone cells. Among the increased structural parameters, the tubule area showed the smallest increase in the cotyledon, at 3.36% compared to the CK group. Conversely, post-RALF treatment, the areal cisternae displayed reductions in all growth parts, particularly in the cotyledon, which was three times greater than that of the hypocotyl and ten times larger than that of the elongation zone, respectively (Figure 2H and 2L).

In the treatment on the 4th day seedling, the area of cisternae in the hypocotyl showed the most significant change, increasing by 99.45% compared to the CK group. However, this parameter exhibited the most substantial decline in the elongation zone (approximately 72.01%). Following the treatment, the growth tip experienced a decrease in the cotyledon and hypocotyl (20.22% and 46.90%, respectively), but it showed an increase in the elongation zone by 7.16%. The number of fusiform bodies, the area of bulk flow, and the area of cisternae and bulk flow exhibited an inverse trend compared to the growth tip; there was an increase in the cotyledon and hypocotyl but a decline in the elongation zone. Notably, the area of bulk flow in the hypocotyl and elongation zone showed the most significant discrepancy (the hypocotyl was 17 times larger than the elongation zone). Moreover, the area and length of tubules, as well as the number of three-way and multi-way junctions, decreased in all three growth parts when compared with the CK group (Figure 2I and 2L). However, on the fifth day, when the seedlings were treated with RALF, almost all changes in structural parameters were less pronounced than in the preceding two days. The number of multi-way junctions in the elongation zone exhibited the most significant change, amounting to 74.38%. The area and length of tubules, along with the number of fusiform bodies, decreased in the cotyledon but increased in the hypocotyl and elongation zone. Additionally, the changes in the area of tubules in the hypocotyl and elongation zone were twice as much as those in the cotyledon (Figure 2J and 2L).

Analyzing data from the 3rd to the 5th days under RALF treatment compared to the CK, we observed distinct variations in all structural parameters across these three growth regions. The cisternae area increased solely in the cotyledon, yet the most substantial change occurred in the elongation zone at 31.97%, approximately 30 times more than the cotyledon and 10 times more than the hypocotyl. Bulk flow area decreased by 1.17% solely in the elongation zone, while increasing by 17.65% and 28.28% in the cotyledon and hypocotyl, respectively. Both in area and tubule length, the decrease observed in the cotyledon and hypocotyl was nearly equivalent to the increase noted in the elongation zone. Post RALF treatment, the number of fusiform bodies increased solely in the hypocotyl by 10.82%. Regarding ER nodes, following RALF treatment, a decrease was observed in both the cotyledon and hypocotyl, while an increase was noted in the elongation zone (Figure 2K and 2L).

Changes in ER structural parameters vary across different growth regions, particularly in response to rising environmental pH levels. Within the cotyledon, characterized by a sheet-like structure such as cisternae and bulk flow, there was an increase observed. Parameters linked to the branched network, including tubules and nodes, exhibited a decrease. Additionally, the count of fusiform bodies decreased within the cotyledon. In the hypocotyl, notable changes primarily manifested in the branched network, resulting in decreased tubules and nodes. Meanwhile, within the elongation zone, alterations predominantly affected the sheet-like structure, while an increase was noted in the branched network.

### Quantitative Analysis of dynamic changing of ER structure in time-lapse

ER is a highly dynamic organelle whose sheet-like structure and branched network undergo changes during reshaping (Lu et al., 2023). To enhance the visual representation of temporal changes in the endoplasmic reticulum (ER) structure, we conducted a three-dimensional analysis of the hypocotyl’s ER on day 3. This analysis encompassed the horizontal x- and y-axes along with the time axis, utilizing 20 time points captured at 30-millisecond intervals during a six-second imaging session. Subsequent imaging at each time point revealed alterations in the ER structure, encompassing tubules, cisternae, and fusiform bodies (Figure 3A). Notably, the cisternae, constituting the predominant component of the sheet-like ER structure, not only contributes significantly to its volume but also undergoes structural modifications upon imaging at each time point (Figure 3B). Meanwhile, as the framework of the branched network in ER, tubule also had changed in 20 time points (Figure 3C). The dynamic movement of fusiform body seemed to be the most obvious in all three structural pamaters. (Figure 3D).

**Figure 3.**
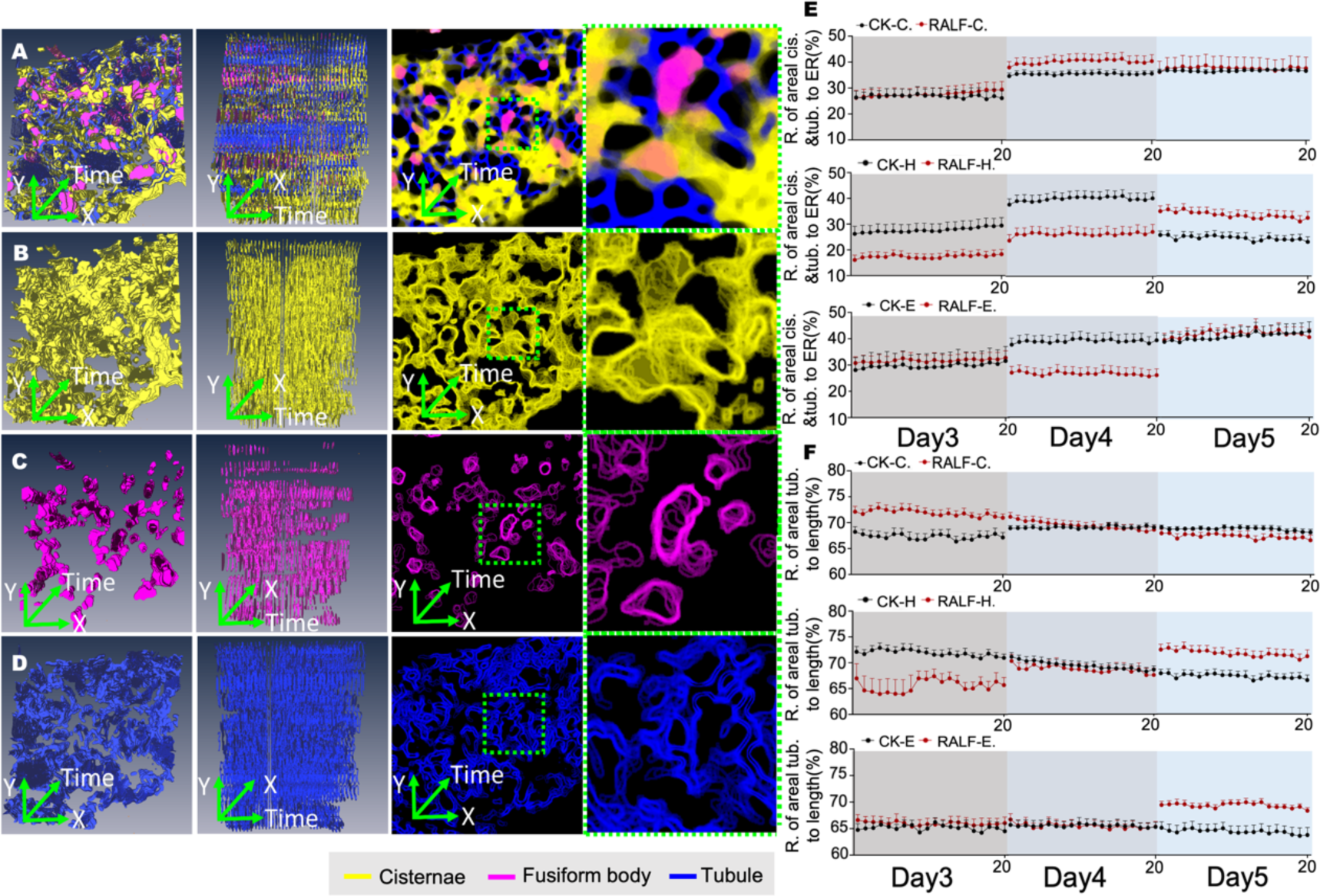
Three-dimensional and time-series analysis. **(A)-(D)** A three-dimensional analysis of the dynamic changing of structure of cisternae, fusiform body, and tubule within time series **(E)** The quantitative analysis of the ratio of areal cisternas and tubule to areal ER in cotyledon, hypocotyl, and elongation zone under RALF treatment with time-lapse **(F)** The quantitative analysis of the ratio of areal tubule to length in cotyledon, hypocotyl, and elongation zone under RALF treatment with time-lapse

Upon analyzing the structural parameters of ER in a 3-day growth period with CK and RALF, we observed evident alterations in the ER structure due to the treatment. There were fluctuations observed in the morphological dynamics of the ER structure across the 20 time points, notably displaying distinctive structural changes under the RALF-induced stress treatment.

The ratio of areal bulk flow to areal ER had changed obviously after the RALF treated on the 3-day-old seedling in cotyledon and hyocotyl. Both the cotyledon and hypocotyl showed an approximate 16% increase in this structural parameter compared to the CK group under the influence of RALF treatment. Nevertheless, the ratio in the RALF group of the hypocotyl (80.71% ± 0.53) exhibited less variability compared to the CK group (67.02% ± 1.13). Furthermore, there appeared to be no discernible impact of the treatment on the fluctuations observed in the cotyledon and elongation zone (Supplemental Figure). After the RALF treatment, the ratio of areal cisternae and tubules in relation to ER showed a decrease in both the cotyledon (from 30.23% ± 0.45 to 19.24% ± 0.38) and hypocotyl (from 32.98% ± 0.53 to 19.29% ± 0.53), yet no significant fluctuation was observed in these three growth regions. Following RALF treatment, there was a reduction in the ratio of areal tubule to length specifically in the hypocotyl (from 71.82% ± 0.48 to 66.66% ± 1.93), with the RALF treatment, this ratio in hypocotyl had the most violent fluctuations among these three growth parts, almost was four times greater than that of the CK group (cotyledon was 68.29% ± 0.18, and the elongation zone was 65.77% ± 0.079). (Figure 3E).

On the 4-day-old seedling treatment, there were notable fluctuations in the ratio of areal bulk flow to ER in both the cotyledon and hypocotyl. In the cotyledon, the treated group exhibited a 75.2% increase in the fluctuation of the ratio (50.80% ± 2.42) compared to the CK group (41.24% ± 0.6). Meanwhile, in the hypocotyl, the change in fluctuation from the CK group (40.58% ± 1.91) to the RALF group (67.30% ± 0.94) increased by 50.79% (Supplemental Figure). Similar fluctuations in the ratio of areal cisternae and tubules to ER were observed in both the cotyledon and hypocotyl (Figure 3E). Following RALF treatment, there was a noticeable fluctuation in the ratio of areal tubule to length across all three growth parts. In the cotyledon and elongation zone, the treated group exhibited an increase of approximately 63% to 67% compared to the CK group. Conversely, in the hypocotyl, the volatility was twice as high in the CK group as in the RALF group (Figure 3F).

Upon treating the 5-day-old seedlings with RALF, noticeable wave-like motions were observed in the ratio of bulk flow to ER within the RALF-treated group. In the cotyledon, the ratio within the RALF group (35.23% ± 1.27) exhibited a 15% increase compared to the CK group (34.15% ± 0.93). Conversely, in the hypocotyl, the RALF group (66.35% ± 0.96) displayed a significant 44.4% increase compared to the CK group (48.71% ± 0.57). Within the elongation zone, although the fluctuations were less pronounced compared to the cotyledon and hypocotyl, a noticeable 10.64% increase was observed post-treatment (Supplemental Figure).

The ratio of areal cisternae and tubules to ER showed consistent fluctuation patterns with the ratio of areal bulk flow to ER in both the cotyledon and elongation zone. Notably, in the hypocotyl, there was a distinct 25% increase in this ratio post-RALF treatment (Figure 3E). Furthermore, there was a substantial 73% increase in the ratio of areal tubule to length specifically in the hypocotyl after treatment (Figure 3F). Post-RALF treatment, significant fluctuations were observed in the ratio of areal bulk flow to ER and the ratio of areal cisternae and tubule to ER across those growth zones, with the most pronounced changes occurring notably in the hypocotyl, particularly on the 4th day. The elongation zone experienced the least impact from RALF treatment. Notably, the most evident fluctuations in the areal ratio of tubules to length continued to be observed in the hypocotyl, while the cotyledon exhibited the least impact from RALF treatment (Figure 3F).

### Analysis ER dynamics in timelapse using optical flow

In Figure 4, we estimate the direction and speed of different changes of sub-compartment of ER from previous optical image frame to the current one using the Horn-Schunck algorithm. Here, we show the dynamic changes of ER compartment by using a image sequence taken from 3-CK-L-11. These sub-compartments of ER including Cisternae Bulkflow, ER Tubule, and FusiformBody have different characteristics of dynamic changes in a measured time period (Figrue 4A-D). We also characterised the dynamic changes of fusiform body (Figure E-H), tubule and cisternae (Figure I-L).

**Figure 4.**
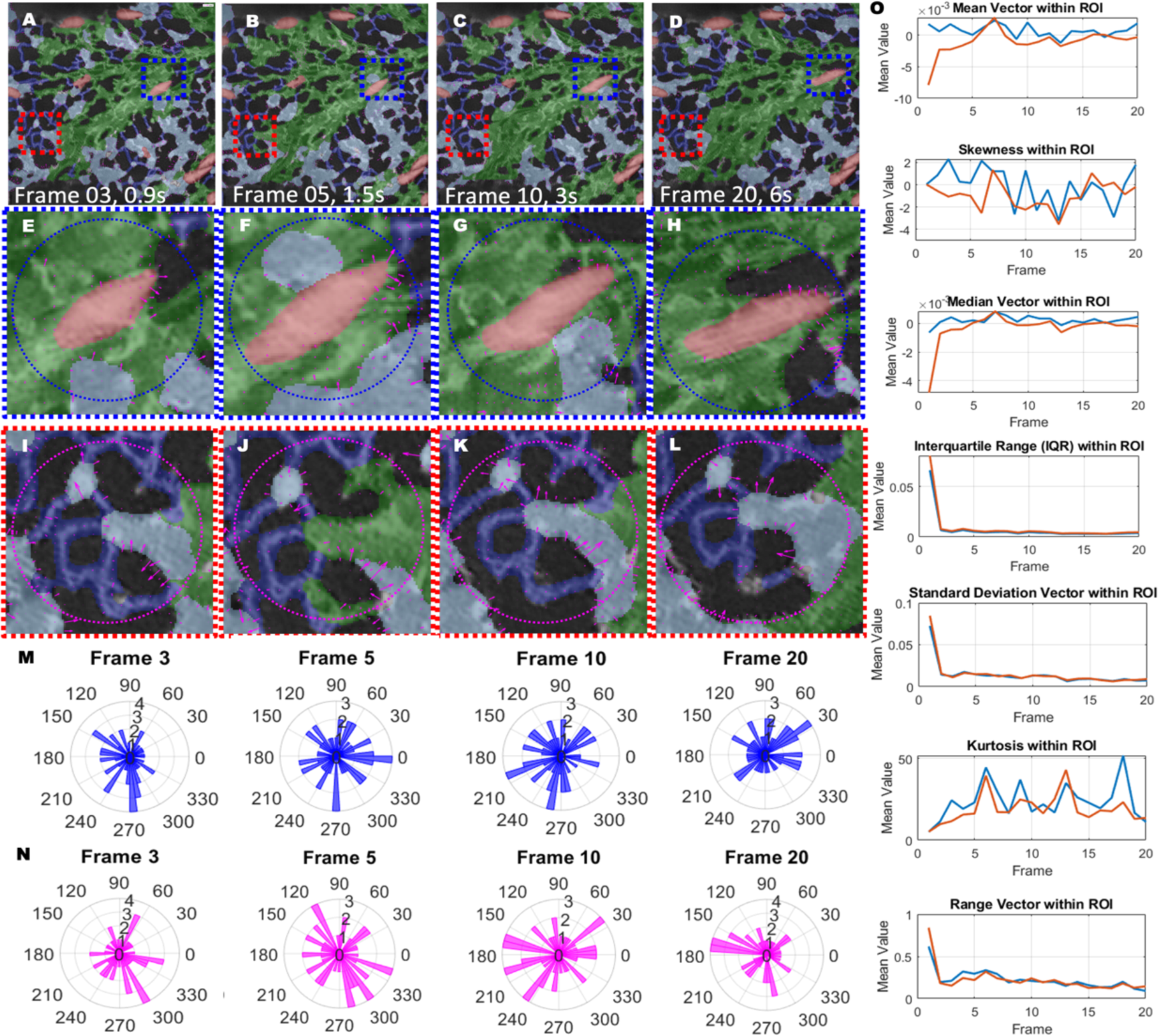
Dynamic changes of different ER compartment using optical flow. **(A)-(D)** Display the identified and segmented sub-compartment of ER in a acquire image sequence (3-CK-L-11). The entire image sequencs has 20 frames taken 6 seconds. **(E)-(H)** Inside region of interest (ROI blue cycle), ROIs show the velocities and directions of dynamic movements and changes of fusiform body in the frame 03, 05, 10, and 20. **(I)-(L)** Inside region of interest (ROI red cycle) , ROIs show the dynamic changes of tubule and cisternae in different timesteps by using the optical flow vectors as quiver plot. **(M)** The vector angle histogram computed from blue ROIs in E-H, the angle histogram of optical flow showing dynamic changes of sub-compartments in ER are different from each frame following the timesteps. **(N)** The vector angle histogram computed from magenta ROIs in I-L, the angle histogram of optical flow showing dynamic changes of sub-compartments in ER are different from each frame following the timesteps.

### Quantitative comparisons using multiple parameters under RALF stress

We analyzed the changing of nodes (growth tip, three-way junction and multi-way junction) in ER and fusiform body - unique structure in ER of *Arabidopsis*, before and after RALF treatment (Supplymantry figure). The number of three-way junction presented a sigficant changing after RALF treatment. For determining those structural data changing, day 3 data was set to 1 for analysis of data changes after RALF treatment on days 4 and 5. On the 4^th^ day, with the RLAF treantment, the ER structure from different growth zone presented differently. The structure parameters of branched network in hypocotyl presented obviously increase. Including the structure of tubule (area and length) increased 56.94% and 40.97% compared with CK group, which were almost twice with those parameters in cotyledon and elonagiton zone (the branched network decreased in both growth parts). Of course, the nodes in hypocotyl increased 4-6 times than cotyledon and elongation zone, especilly for the three-way junction and multi-way junction which are created by the branched network presented an oppsite tend than others (the three-way jucntion in hypocotyl increased 60.94% and decreased 111.08% and 196.4% in cotyledon and elongation zone). On the 5^th^ day, the same changing tends happened in all of thoes three growth parts after RALF treatment (Figure 5A,5B). The branched network in ER from those three growth parts preseneted an increased tend after RALF treatment, especially in hypocotyl. However, the flat membrane network in cotyledon also increased but not as much as the branched network.

**Figure 5.**
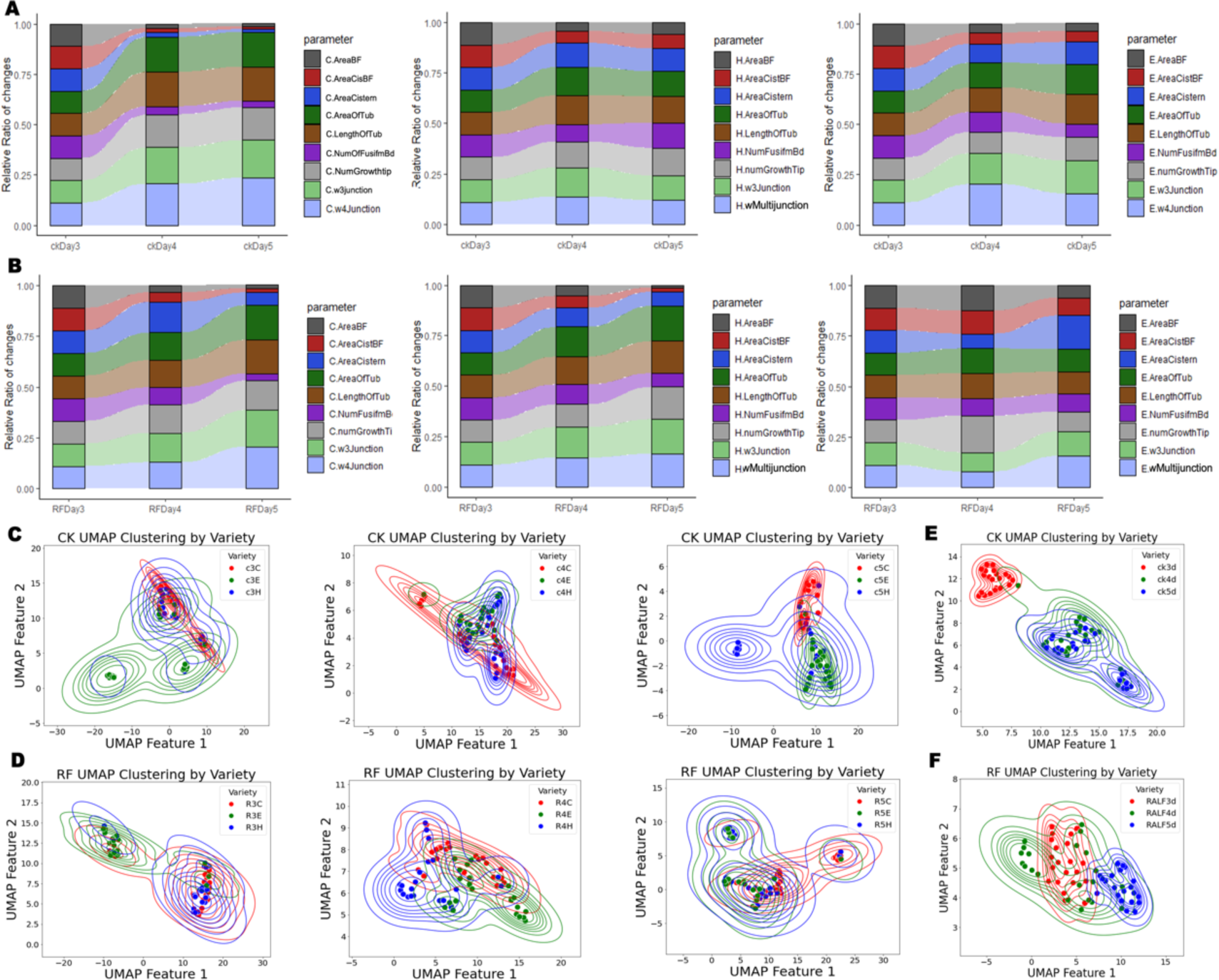
Data distribution and UMAP cluster comparative analysis. **(A)-(C)** Nine parpamters of ER structure changes in 3-5 growth day under CK group **(D)-(F)** Nine parpamters of ER structure changes in 3-5 growth day under RALF treatment **(G)** UMAP clustering analysis with CK group during 3-5 growth day **(H)** Comprehensive UMAP clustering analysis with CK group within all growth day **(I)** UMAP clustering analysis with RALF treatment during 3-5 growth day **(J)** Comprehensive UMAP clustering analysis with RALF treatment within all growth day

**Figure 6.**
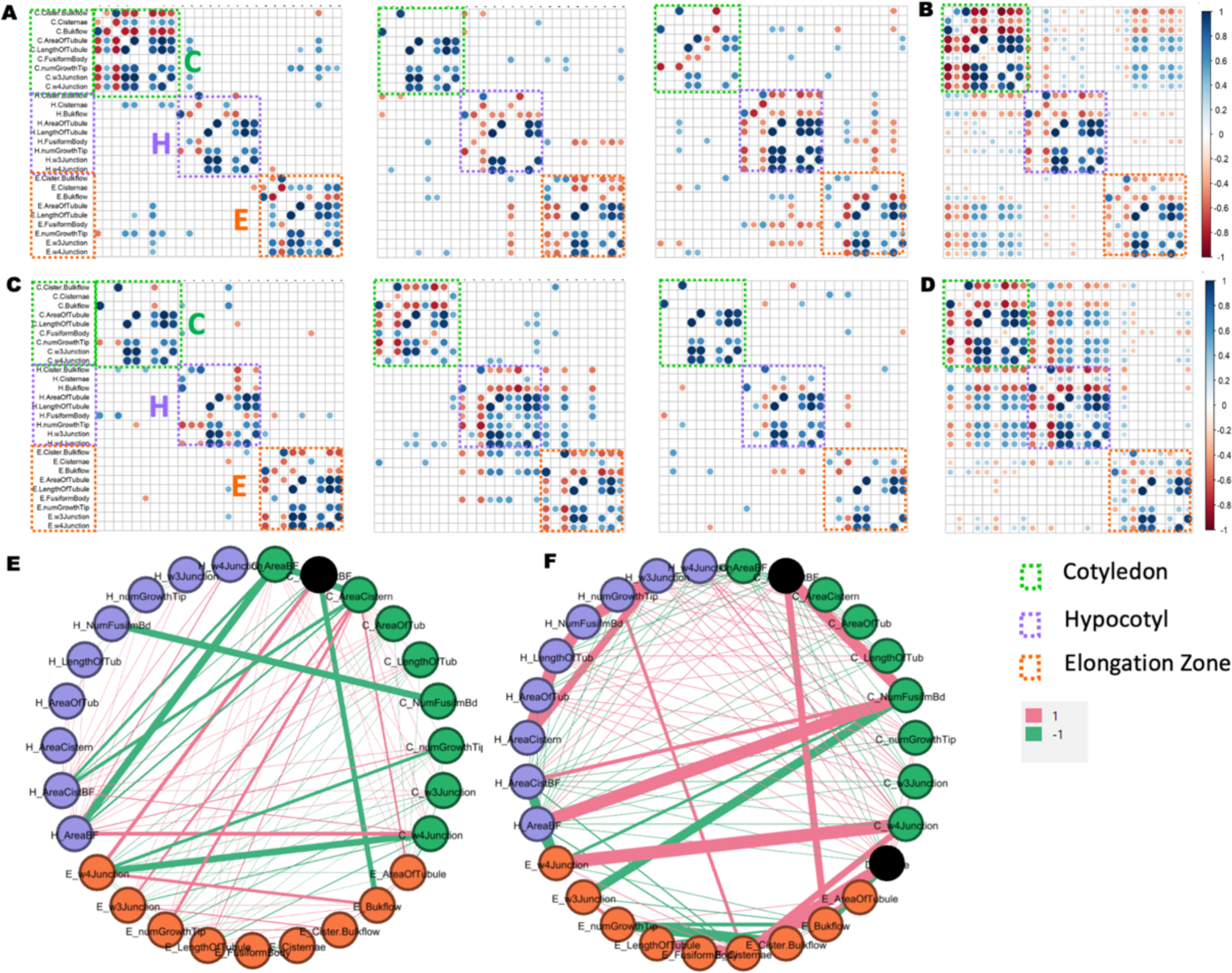
Correlation of changes with respect to CK and RALF. **(A)** The pearson correlation matrix of Cotyledon and Hypocotyl and Elongation Zone of CK acquired are plotted in scatter plots with respect to Day 3, day 4 and day 5 **(B)** the scatter plot of The pearson correlation matrix of Cotyledon and Hypocotyl and Elongation Zone of CK based on the combination of day3, day4 and day5 **(C)** The pearson correlation matrix of Cotyledon and Hypocotyl and Elongation Zone of RALF acquired are plotted in scatter plots with respect to Day 3, day 4 and day 5. **(D)** the scatter plot of The pearson correlation matrix of Cotyledon and Hypocotyl and Elongation Zone of RALF based on the combination of day3, day4 and day5. In figure, 1 denoted positive correlation and -1 denoted negative correlation between each node.

Uniform Manifold Approximation and Projection (UMAP) which is a robust dimensionality reduction and visualization technique extensively utilized in data analysis and machine learning. It effectively maps high-dimensional data onto a lower-dimensional space, preserving the local and global structures of data points. In our study concerning mass ER structural patamars data, UMAP provided visual insights into nine primary ER structural patamars across three growth parts under varying RALF stress conditions. Upon analyzing the distribution of these clusters post RALF treatment in 3-day-old seedlings, both cotyledon and hypocotyl clusters appeared to occupy similar regions in the UMAP. However, subsequent to RALF stress, the elongation zone cluster demonstrated increased centralization. Following the RALF treatment in 4-day-old seedlings, all three clusters exhibited decentralization. As for the RALF treatment in 5-day-old seedlings, all clusters, except for a minor portion of elongation zone parameters, displayed increased centralization (Figure 4C, 4D).

Upon summarizing data spanning days 3 to 5 post-RALF treatment, the cotyledon cluster demonstrated increased proximity with the other two clusters (hypocotyl and elongation zone). Conversely, these two clusters exhibited decentralization tendencies after RALF treament (Figure 4E, 4F)

### Correlation analysis under RALF stress

Under stress conditions, the structural parameters of the endoplasmic reticulum in different parts of cells are different.The trifurcation point in the leaves is most obviously reduced under stress conditions. Overall flow increases under stress conditions in three different cells In three different cells, except for the elongation zone, the area ratio of tubes decreased under stress conditions, which was more obvious in leaves and hypocotyls. The spindle body of cells in the elongation zone decreases most significantly under stress conditions.The right column showed the scatter plots and the corresponding linear Pearson correlation coefficient (rp) and the nonlinear Spearman correlation coefficient (rs), indicating the extent of colocalization with the value less than zero corresponds to a negative relationship. Positive correlation +1, and negative correlation -1. Under stress conditions, the structural parameters of the endoplasmic reticulum in different parts of cells are different.

## DISCUSSION

RALF has many receptors, like CrRLK1L, can form complexes that involved in many aspects of plant development and growth, reproduction, hormone signaling, immunology, and stress responses (Yu et al., 2012; Stegmann et al., 2017; Song et al., 2022; He et al., 2023). Those receptors distributed in all plants except pollen. Most receptors are located in the root to perform physiological functions, like

## METHODS

### Plant Materials and Growth Conditions

The plant material used in this experiment was col-0 ecotype *Arabidopsis thaliana* expressing the HDEL-GFP fluorescent tag. *Arabidopsis* seeds were surface sterilized for 30 s in mixed sterilized water (75% ethanol and 5% H_2_O_2_) and stratified at 4°C for 48h in darkness. The seedlings were then grown on vertical plates containing half-strength Murashige and Skoog (MS) medium (pH=5.95) in a growth chamber with a constant temperature of 22°C, relative humidity of about 70% and 16 h light/8 h dark cycles. Seedlings grown for 3 days, 4 days and 5 days were taken for imaging observation as per the experimental requirements.

### Stress Treatments

The RALF peptides were synthesized by SciLight Biotechnology and used at a concentration of 1 µM in half-strength MS liquid growth medium with the same pH as the solid growth medium. *Arabidopsis* seedlings were treated by immersion in RALF solution for 30 min to simulate alkaline stress conditions. The control (CK) group was liquid 1/2MS without RALF addition.

### Denosie-enhance Framework and Segmentation

The processing programs in this paper are based on python platform. Firstly, the Blind2Unblind implicit lossless self-supervised denoising framework is used for image denoising and enhancement; subsequently, the U-net based segmentation network is applied for image recognition.

Leveraging the segmentation outcomes derived from Swint ResU-Net, we engineered a graph incorporating elements such as Tubule, cisternae, bulk flow, and the fusiform body. We pinpointed consistent nodes within this network, which included the growth tip, three-way junction, and multi-way junction.

### SIM Microscopy

SIM fluorescent images were collected using a Deltavision OMX structured light illumination super-resolution microscope (model: deltavision OMX SR) with 60x oil lens. Arabidopsis seedlings were taken to make a clinical slide and placed on the carrier stage, the fluorescence signal for GFP (488nm), the laser intensity was 15%, the exposure time was 30 ms, the TIRF-SIM has three angles and its si tirf pitch was 4.45 (0.00, 0.00, 0.00), and 20 consecutive frames of images were captured in a total of 6s. The Fourier algorithm was used to reconstruct and generate the super-resolution imaging of the time series.

## Author Contribution

Yiheng Zhang acquired and anaylzed the data. Jiazheng Liu prepared the denosie framework and the Swint ResU-Net for segmentation ER structure and prepared the figure of Swint ResU-Net algorithm. Yiheng Zhang prepared all figures based on segmented image data made by Liu.

## Funding

STI 2030-Major Projects (2021ZD0204500, 2021ZD0204503 to L.L.), National Natural Science Foundation of China (No. 91954202, 31871349 to X.L. No. 32171461 to H.H.,), Scientific research instrument and equipment development project of Chinese Academy of Sciences (YJKYYQ20210022 to H.H.), The Third Xinjiang Scientific Expedition Program (Grant No.2022xjkk1200), Beijing Forestry University Outstanding Young Talent Cultivation Project (2019JQ03003)

## Notes

### Competing Interest Statement

The authors have declared no competing interest.

